# Crosslinking of nucleotide binding domains improves the coupling efficiency of an ABC transporter

**DOI:** 10.1101/836676

**Authors:** Chengcheng Fan, Jens T. Kaiser, Douglas C. Rees

## Abstract

ATP Binding Cassette (ABC) transporters often exhibit significant basal ATPase activity in the absence of transported substrates. To investigate the factors that contribute to this inefficient coupling of ATP hydrolysis to transport, we characterized the structures and functions of variants of the bacterial Atm1 homolog from *Novosphingobium aromaticivorans* (*Na*Atm1), including forms with disulfide crosslinks between the nucleotide binding domains. Unexpectedly, disulfide crosslinked variants of *Na*Atm1 reconstituted into proteoliposomes not only transported oxidized glutathione, but also exhibited more efficient coupling of ATP hydrolysis to GSSG transport than the native transporter. These observations suggest that enhanced conformational dynamics of reconstituted *Na*Atm1 may contribute to the inefficient use of ATP. Understanding the origins of this uncoupled ATPase activity, and reducing the impact through disulfide crosslinking or other protocols, will be critical for the detailed dissection of ABC transporter mechanism to assure that the ATP dependent steps are indeed relevant to substrate translocation.

## Introduction

Membrane transporters couple the translocation of ligands to a thermodynamically favorable driving force. A key feature of the transport mechanism is that these two processes - ligand transport and the driving force - must be linked to minimize “short circuiting” of the transduction process (Tanford 1983, Hill 1989). In essence, transporters kinetically facilitate the coupled reaction while disfavoring the individual uncoupled reactions. One way to achieve coupling between the favored and unfavored reactions is when both processes proceed through a common set of conformational states so that they can only be accessed when the relevant components are both present (Rees and Howard 1999). Structure based mechanisms defining these coupling processes have been advanced for several transporters, including lac permease (Kaback 2015) and P-type ATPases (Palmgren and Nissen 2011).

In contrast to those systems, ATP Binding Cassette (ABC) transporters provide an interesting situation where the coupling mechanism is less well defined. The transport mechanism is generally described by the alternating access model involving inward-facing, occluded and outward-facing states, with the transitions between states coupled to the binding and hydrolysis of ATP, followed by product release (Hofmann et al. 2019). A striking feature of certain characterized ABC transporters, including exporters and importers (Lewinson and Livnat-Levanon 2017), is the significant uncoupled ATPase activity in the absence of substrate, reflected in high basal ATPase activities, and the inefficient coupling of ATP hydrolysis to substrate transport, reflected in the ratio of hydrolyzed ATP to translocated substrate (Table 1). While the general view of ABC transporters is that the coupling ratio is ~2 ATP per transported substrate, in practice, ratios near this value have been reported for only a few transporters and more typically greatly exceed 2. Understanding and overcoming the underlying causes of the generally poor coupling efficiencies is critical for advancing the quantitative mechanistic understanding of ABC transporters.

**Table 1.** Coupling efficiencies between ATP hydrolysis and substrate translocation for ABC transporters. Coupling efficiencies are either presented in the corresponding reference or calculated based on the reported ATPase and transport activities of the transporter. Coupling efficiency = ATPase activity/transport activity.

| Transporters | ATPase activity (nmol/min/mg) | Transport Activity (nmol/min/mg) | Coupling efficiency | References |
| --- | --- | --- | --- | --- |
| **ABCC3** | 200 | 1,200 | 0.17 | Zehnpfennig et al. 2009 |
| **MalFGK** | ~3 | ~2 | 1.4 - 17 | Davidson et al. 1990 |
| **OpuA** | ~80 - 120 | ~30 - 70 | 1.7 - 4 | Patzlaff et al. 2003 |
| **GlnPQ** | 15 (min<sup>-1</sup>) | 8.5 (min<sup>-1</sup>) | 1.8 | Lycklama et al. 2018 |
| **Pgp** | ~750 - 1,300 | ~500 | 2 | Eytan et al. 1996 |
| **ABCG5/8** | 110 | 50 | 2.2 | Wang et al. 2006 |
| **MalFGK** | ~1.2 - 8 | ~0.3 - 2 | 4 - 10 | Dean et al. 1989 |
| **Pgp** | ~110 | ~6 | 18 | Dong et al. 1996 |
| **HisP** | 580 | 19 | 31 | Nikaido and Ames 1999 |
| **ABCG2** | ~750 | ~22 | 34 | Manolaridis et al. 2018 |
| **TmrAB** | ~1,100 | ~30 | 37 | Hofmann et al. 2019 |
| **BtuCDF** | ~400 | ~4 | 100 | Borths et al. 2005 |
| **HmuUV** | ~130 | ~1.1 | 120 | Woo et al. 2012 |
| **NaAtm1** | 180 | 1.5 | 120 | This study |
| **MalFGK** | 4,000 | 1.2 | 3,300 | Chen et al. 2001 |
| **ABCB6** | 610 | 0.03 | 20,000 | Chavan et al. 2013 |

To investigate the factors that contribute to the high basal ATPase activity and coupling inefficiencies of ABC transporters, we characterized the structures and transport functions of variants of a bacterial homolog of the ABC transporter of mitochondria (Atm1) from *Novosphingobium aromaticavorans* (*Na*Atm1). Atm1 is a homodimeric exporter that was first reported to be involved in iron-sulfur cluster biogenesis in yeast (Kispal et al. 1997). The human homologs of Atm1, ABCB6 and ABCB7 (Lill and Kispal 2001), have been implicated in iron homeostasis (Kelter et al. 2007, Cavadini et al. 2007, Pondarre et al. 2006, Kispal et al. 1997); the plant homolog, Atm3 from *Arabidopsis*, is important for both iron-sulfur cluster biogenesis (Bernard et al. 2009, Zuo et al. 2017) and molybdenum cofactor maturation (Teschner et al. 2010). While the physiological substrates for these transporters are not well defined, earlier studies (Kuhnke et al. 2006) suggested that glutathione (GSH, reduced; GSSG, oxidized forms) and derivatives, including persulfidated forms (Schaedler et al. 2014, Riedel et al. 2019), are the substrates for Atm1 homologues. Our initial structural and functional studies of the bacterial homolog support a role of Atm1 in in heavy metal detoxification, plausibly by export of metallated glutathione species (Lee et al. 2014).

In this report, we characterize the GSSG transport and ATPase activities of wildtype *Na*Atm1 and variants with disulfide crosslinked NBDs. Unexpectedly, a disulfide stabilized variant exhibited more efficient coupling between ATP hydrolysis and transport, while retaining a transport rate 60% of the wild type transporter. These findings suggest that the enhanced conformational dynamics of reconstituted *Na*Atm1 may contribute to the high basal ATPase rate and the inefficient coupling to substrate transport.

## Results

### Disulfide bond crosslinking

The conformational state of an ABC transporter is reflected in the arrangement of the NBDs which can vary from well-separated in the inward-facing conformation to a fully dimerized state in the outward facing form. As an approach to addressing how the accessibility of different conformational states influences the transport cycle of *Na*Atm1, we introduced disulfide bridges at the dimerization interface between the NBDs to stabilize the transporter in conformational states with juxtaposed NBDs (Korkhov, Mireku, and Locher 2012). Through sequence and structural alignments of *Na*Atm1 to ABC transporters with dimerized NBD structures (Figure S1a), stabilized either by nucleotide binding (Dawson and Locher 2006) or a disulfide bridge (Korkhov, Mireku, and Locher 2012), we identified three residues, A527, S526 and T525, near the Walker-B motif where disulfide bonds could potentially form between the equivalent residues in the two NBDs following substitution with cysteine and oxidation. These three residues were separately mutated to cysteine in the natively cysteine-less homodimeric *Na*Atm1 to generate three single-site variants: *Na*A527C, *Na*S526C, and *Na*T525C. All variants exhibited similar crosslinking yields in the initial crosslinking tests (Figure S1b). The availability of these disulfide crosslinked variants provided an approach to address the functional and structural properties of *Na*Atm1 upon constraining the relative positions of the NBDs.

### Proteoliposome reconstitution

To test the ATPase and transport functions of *Na*Atm1 in a membrane like environment, we separately reconstituted into proteoliposomes (PLS) wildtype *Na*Atm1, the cysteine variants (*Na*A527C, *Na*S526C, and *Na*T525C) and *Na*E523Q, the ATP-hydrolysis deficient variant with the E to Q mutation in the Walker B-motif (Moody et al. 2002) by following an established protocol (Geertsma et al. 2008). The incorporation efficiency of *Na*Atm1 into PLS was evaluated by running samples on SDS-PAGE of the total PLS and the supernatant following ultracentrifugation. With a final reconstitution efficiency above 95% (Figure S2a), it was assumed full incorporation for subsequent analyses. Transporters reconstituted into PLS can adopt two possible orientations with the NBDs positioned either outside or inside of the liposomes. In these studies, we utilized transporters oriented with the NBDs on the outside to measure both the ATPase and the transport activities (Figure 1a). In contrast, only the ATPase activities could be measured for *Na*Atm1 purified in detergent (Figure 1b). Following collection of the PLSs, transport activities were measured with a glutathione reductase based enzymatic assay, and the ATPase activities were measured using a molybdate based colorimetric assay (Chifflet et al. 1988).

**Figure 1.**
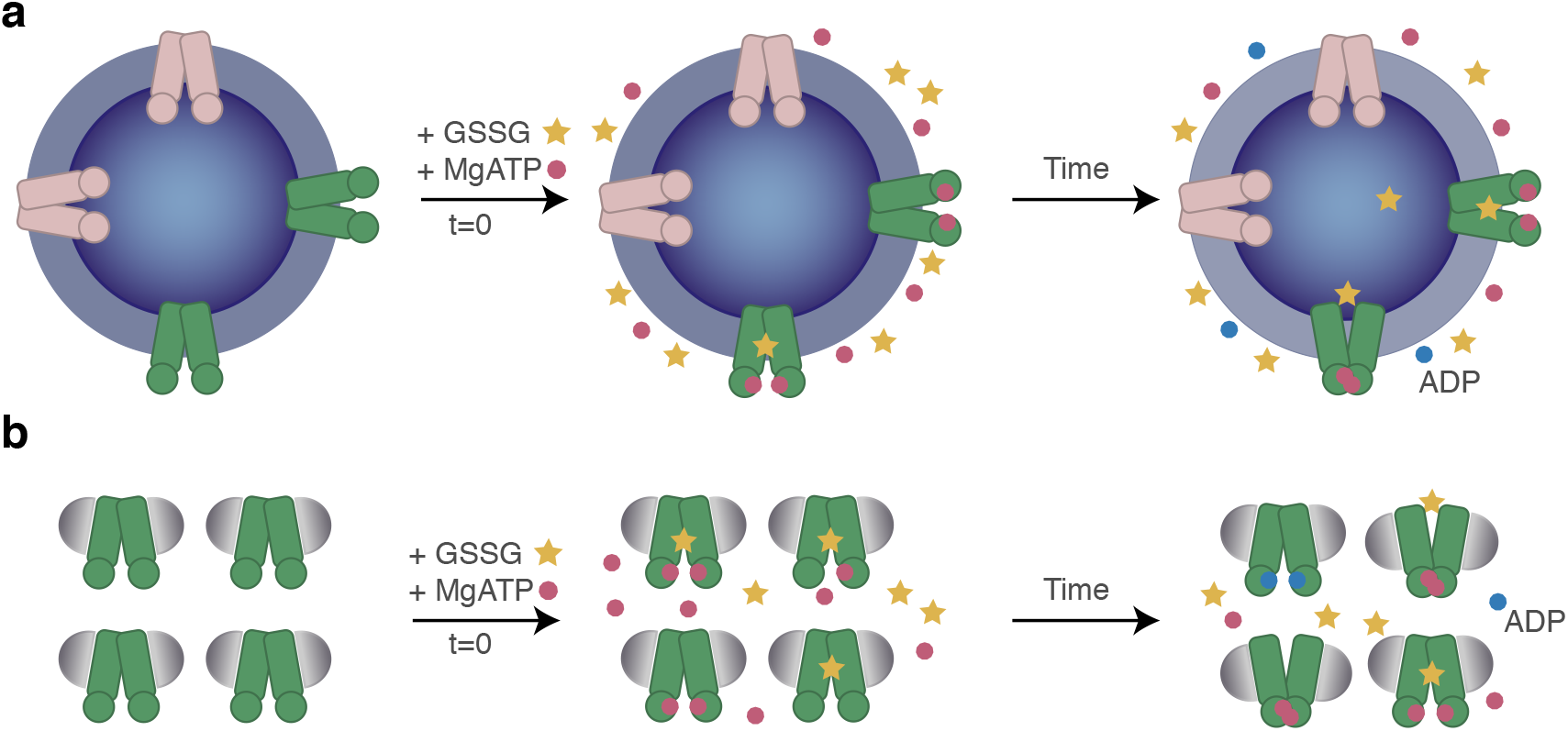
Functional assay schematics. (**a**) Transport and ATPase assay in proteoliposomes illustrating the two possible orientations of a transporter reconstituted in proteoliposomes. (**b**) ATPase assay in detergent.

### Transport and ATPase activities of wildtype and *Na*Atm1 mutants

We first established the transport assay with wildtype *Na*Atm1 reconstituted in PLS, with MgATP and GSSG present at physiological concentrations, 10 mM and 2.5 mM, respectively (Bennett et al. 2009) on the outside of the PLS. Transport activity was quantitated by recovering the PLS and measuring the accumulation over time of GSSG inside the PLS (Figure S2b). The negative controls showed no measurable time dependent GSSG uptake, although it appeared that low levels of GSSG stick to PLS or liposomes. For wildtype *Na*Atm1, the uptake of GSSG was approximately linear with time, corresponding to a rate of 1.5 nmol GSSG min^−1^ mg^−1^ transporter (Figure 2a, S2b). Based on the *Na*Atm1 molecular weight of 133 kDa (with 1 mg = 7.5 nmole), neglecting orientation effects and assuming all the transporters are functionally active, this rate is equivalent to ~0.2 GSSG translocated per minute per transporter.

**Figure 2.**
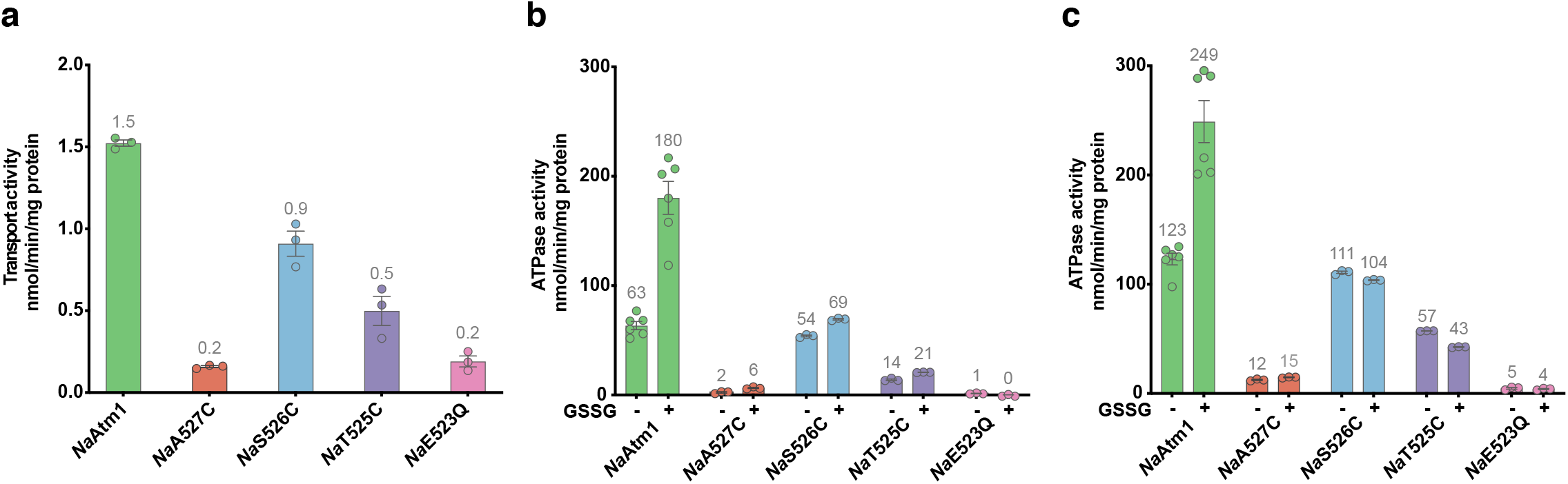
*Na* Atm1 transport and ATPase activities. (**a**) Transport activity of *Na*Atm1 and its variants with 10 mM MgATP and 2.5 mM GSSG. (**b, c**) ATPase activities of *Na*Atm1 and its variants at 10 mM MgATP in the absence and presence of 2.5 mM GSSG, in PLS (**b**) and detergent (**c**). Error bars represent the standard error of the mean. Circles in **a, b** and **c** represents the resuls of individual measurements.

Additionally, we measured the transport and ATPase activities of the different mutants for comparison to wildtype *Na*Atm1 (Figure 2). The wildtype *Na*Atm1 reconstituted in PLS showed an ATPase activity of ~60 nmol Pi min^−1^ mg^−1^ transporter with about 3-fold stimulation in the presence of 2.5 mM GSSG to ~180 nmol Pi min^−1^ mg^−1^ transporter (Figure 2b); with *Na*Atm1 purified in detergent, the basal ATPase activity was ~120 nmol Pi min^−1^ mg^−1^ transporter and 2.5 mM GSSG only stimulated the ATPase activity by 2-fold to ~250 nmol Pi min^−1^ mg^−1^ transporter (Figure 2c). *Na*A527C showed significantly reduced transport and ATPase activities in both PLS and detergent (Figure 2). Interestingly, *Na*S526C showed about 60% of wildtype transport activity at a rate of 0.9 nmol GSSG min^−1^ mg^−1^ transporter (Figure 2a), with a basal ATPase activity in PLS similar to wildtype with slight stimulation by GSSG (Figure 2b). *Na*T525C retained about 30% of wildtype transport activity with reduced ATPase activities in PLS and detergent (Figure 2). Lastly, *Na*E523Q exhibited little ATPase activities (Figure 2bc). Both *Na*A527C and *Na*E523Q exhibited a GSSG uptake rate about ~10% of the wildtype protein transport activity despite their diminished ATPase activities (Figure 2a), which may reflect binding of substrate to the transporters.

### Inward-facing occluded conformations

The structures of the different variants were determined by X-ray crystallography to assess the consequences of the disulfide crosslinks on the conformational state of *Na*Atm1. The crosslinked *Na*A527C form with bound MgADP crystallized in space group P1 with four transporters per asymmetric unit (Figure S3a). The initial crystallization trials were carried out with *Na*A527C containing ATP, but the same crystals were obtained with ADP in the crystallization conditions, suggesting these MgADP bound conformations could be achieved either by the slow hydrolysis of ATP or the direct binding of ADP. While each transporter adopted an inward-facing occluded conformation, two distinguishable states (#1 and #2) were evident. Three of the transporters were found in state #1 (Figure 3a), while the fourth transporter in state #2 exhibited a slightly more closed NBD dimer (Figure 3b). Given the moderate 3.7 Å resolution and anisotropic diffraction, we confirmed the two distinct conformations from the locations of selenium sites in selenomethionine substituted protein crystals (Figure S3b). Clear electron density for the disulfide bridges was present in all four transporters (Figure S3c). The primary difference between these structures is reflected by a relative rotation of the NBD α-helical subdomains about the molecular two-fold axis of the transporter (Figure S4a).The root-mean-square deviation (RMSD) between states #1 and #2 is 1.7 Å, while the RMSDs to the previous determined inward-facing structure of *Na*Atm1 are 2.1 Å and 4.4 Å, respectively (Figure S4bcd).

**Figure 3.**
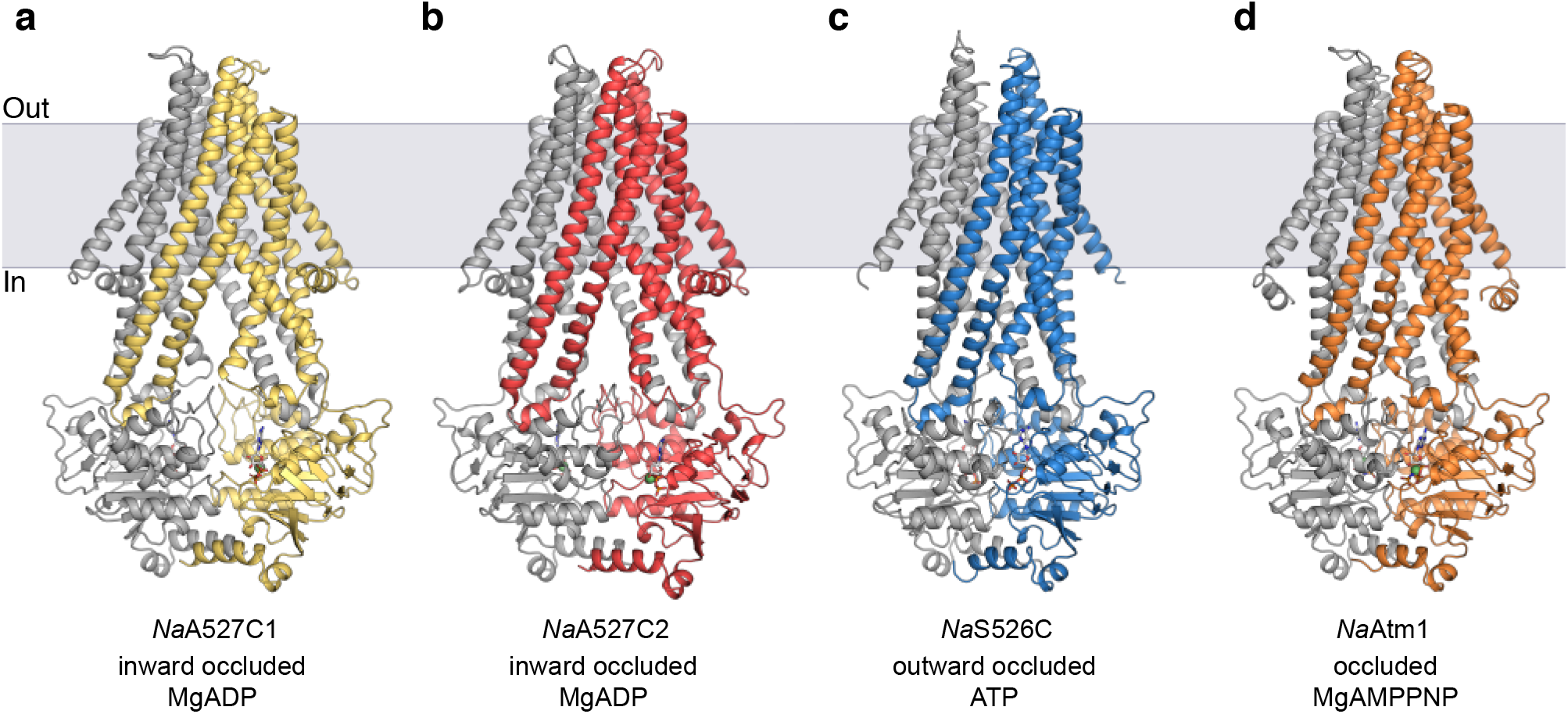
Crystal structures of *Na* Atm1. (**a,b**) Structures of *Na*A527C in the inward-facing occluded conformation #1 (**a**), and #2 (**b**), both with MgADP bound. (**c**) Structure of *Na*S526C in the outward-facing occluded conformation with ATP bound. (**d**) Structure of wildtype *Na*Atm1 in a fully occluded conformation with MgAMPPNP bound. Each of the structures is colored with one chain in gray and the second chain in a different color. The corresponding nucleotides are shown in sticks with Mg^2+^ as green spheres.

Although GSSG was present in the crystallization conditions, ordered electron density for GSSG was not observed in the binding site. To assess whether this ligand was present, we co-crystallized *Na*A527C with a glutathione-mercury complex (GS-Hg) and collected data at the Hg absorption edge. Anomalous electron density peaks identifying Hg sites were found in all GSSG binding sites and were strongest in the two best resolved transporters in the asymmetric unit (Figure S4e). These sites were not observed in control studies using mercury compounds in the absence of glutathione, supporting the interpretation of GS-Hg complex presence in the binding site, which also reflects the binding capability of substrate at the binding site of *Na*A527C.

### Outward-facing occluded and occluded conformations

*Na*S526C, *Na*T525C and *Na*E523Q were each crystallized by removing MgCl_2_ from the crystallization condition of *Na*A527C, followed by minor optimizations. The subsequent structure determinations revealed that these variants adopted similar ATP bound outward-facing occluded conformations (Figure 3c, S5ab). While there is clear electron density for the disulfide bond in the crystal structure of *Na*S526C (Figure S5c), the positions of the cysteine residues in *Na*T525C (separated by 13 Å) are incompatible with the presence of a disulfide bridge in the crystal structure of this variant (Figure S5d). Since no reductant was added, presumably the uncrosslinked population of *Na*T525C was crystallized. The RMSDs between these different outward-occluded structures are ~0.5 Å (Figure S5efg). As the periplasmic regions (residues 60-82 and 284-300) were not well resolved, likely due to the lack of stabilizing crystal lattice contacts, these loops were modeled based on previous structures and refined with low occupancies. Despite the presence of GSSG or GS-Hg in the crystallization conditions, no evidence for substrate binding was observed.

In addition to these mutants, we were able to crystallize and determine the crystal structure of a fully occluded state of wild type *Na*Atm1 with MgAMPPNP bound at 3.35 Å resolution (Figure 3d). This occluded structure shared similar overall architecture as the other outward-facing occluded structures, with alignment RMSDs of ~1 Å (Figure S6). The main distinction is in the periplasmic loop regions that were fully resolved in this electron density map.

## Discussion

Within the framework of the alternating access mechanism, the transition between inward- and outward-facing conformations of ABC transporters proceeds through various occluded states coupled to the binding and hydrolysis of ATP, followed by product release. A recent analysis of the TmrAB heterodimeric drug export established that the outward-facing conformation is stabilized by the binding of MgATP or the MgADP-Pi hydrolysis products, while Pi dissociation accompanies the return to the inward facing conformation (Hofmann et al. 2019). In this study, we have expanded the structurally characterized conformations of the ABC exporter *Na*Atm1 from the initially determined inward-facing conformation to multiple occluded conformations through the use of disulfide-crosslinking of the NBDs and different nucleotides. As observed for previously characterized ABC transporters, different conformations are associated with different ligands. For *Na*Atm1, we have observed that both the outward-facing occluded state and the fully occluded state are stabilized by binding of ATP, or the analog, MgAMPPNP, while the inward-facing and the inward-facing occluded conformations have been observed either nucleotide-free or with bound MgADP. The substrate GSSG has only been observed to bind to inward-facing conformations (Lee et al. 2014), at least at concentrations up to 5 mM.

Previous structural alignments of available exporters structures using a single subunit established that the major conformational differences between various states involve the movement of TM4 and TM5 toward the translocation pathway as the transporter adopts an outward-facing conformation (Lee et al. 2014). With structural alignment of all the *Na*Atm1 structures, changes in TM6 helices between the different conformations are also evident (Figure 4a). In the inward-facing and inward-facing occluded structures of *Na*Atm1, the TM6 helices are kinked, in contrast to the rather straight TM6 helices in the occluded structure and the more subtly bent TM6 helices in the outward-occluded and occluded structures of *Na*Atm1 and the outward-facing Sav1866 (Dawson and Locher 2006). The kinks in TM6 helices associated with the inward-facing conformation occur near two methionine residues, Met317 and Met320, that were previously noted to be involved in the binding of substrates in the inward-facing conformation (Lee et al. 2014). The bends in the TM6 helices observed in the outward-occluded conformations have shifted towards the N-terminal residues by roughly a helical turn to residue Arg313, suggesting that the conformation of TM6 may be sensitive to the presence or absence of bound substrate by *Na*Atm1. These changes in TM6 differ from observations on ABCB1 where substrate binding is accompanied by kinks in the TM4 and TM10 helices (Alam et al. 2019, Alam et al. 2018).

**Figure 4.**
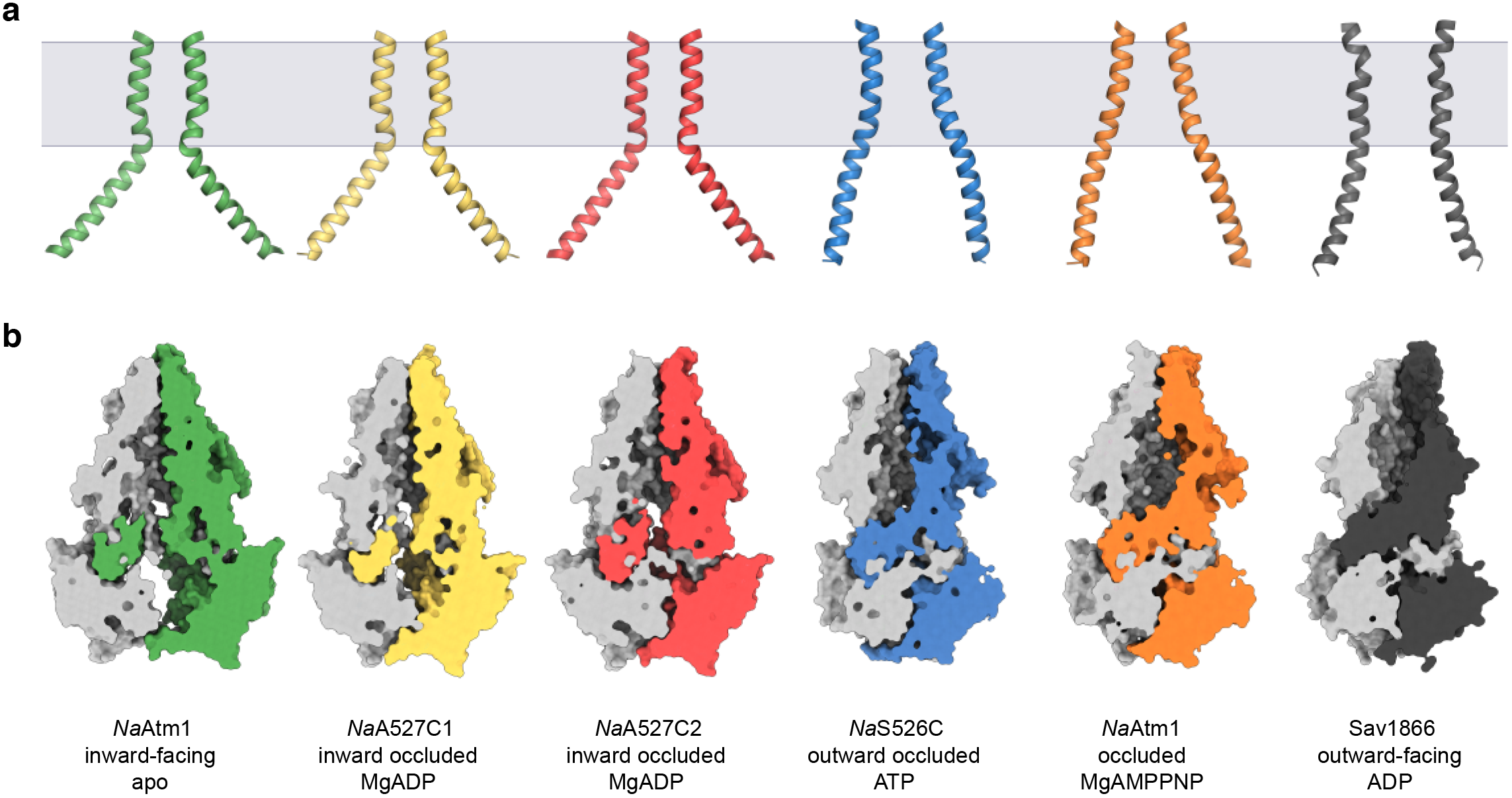
Structural comparisons of *Na* Atm1 structures. (**a**) TM6 (residues 300-340) arrangements of all *Na*Atm1 structures in comparison to Sav1866 (residues 280-320). (**b**) Surface representations of the *Na*Atm1 structures in comparision to our earlier inward-facing structure (PDB ID: 4MRN) and the structure of Sav1866 in the outward-facing conformation (PDB ID: 2HYD) with a slice through the middle of the protein showing the sizes of the central cavities and possible substrate exit tunnels. *Na*Atm1 inward-facing structure (PDB ID:4MRN) in green, *Na*A527C inward-occluded structure #1 in yellow, *Na*A527C inward-occluded structure #2 in red, *Na*S526C outward-occluded structure in blue, *Na*Atm1 occluded structure in orange and Sav1866 outward-facing structure (PDB ID: 2HYD) in black.

The different conformational states of *Na*Atm1 are associated with changes in the substrate binding cavities (Figure 4b). The sizes of the cavities in the inward-facing occluded structures of *Na*A527C are similar to the original inward-facing structure, with sufficient space for substrate binding, as suggested by the GS-Hg anomalous maps (Figure S4e). In the occluded structure of *Na*Atm1 stabilized with MgAMPPNP, the periplasmic gate is fully closed. The three outward-facing occluded structures, represented by *Na*S526C, present slim access paths from the periplasmic end, but the pathways were not as wide open as in the Sav1866 outward-facing structure (Dawson and Locher 2006) (Figure 4b). Although the putative substrate binding cavities in the fully occluded and outward-occluded structures are of sufficient volume to accommodate GSSG (calculated as 1100-1500 Å^3^ with CastP using a 3 Å probe radius (Tian et al. 2018)), the addition of GSSG did not result in extra electron density in the substrate binding sites in the crystallographic studies. These observations suggest these structures may represent the post-translocation states of the transporter.

To assess the functional competence of the constructs generated for these studies, we measured the transport and ATPase activities for *Na*Atm1 and its variants. The activities were determined under physiological concentrations of MgATP and GSSG (10 mM and 2.5 mM, respectively (Bennett et al. 2009)), and the observed values are within the range of values observed for other ABC transporters (Table 1). As a measure of coupling efficiency, we calculated the number of ATP hydrolyzed per translocated GSSG. This calculation may be performed in two ways (Table 2): (i) by taking the ratio of the total ATPase rate to the transport rate (“Total/transport” row in Table 2), or (ii) by taking the ratio of the stimulated ATPase rate (i.e. the additional ATPase rate in the presence of GSSG) to the transport rate (“Stimulated/transport” in Table 2). By focusing only on the increase in ATPase rate in the presence of transport substrate, the latter quantity presumably more accurately reflects the ATPase rate coupled to transport. Either way, it is apparent that the coupling of ATP hydrolysis to substrate transport, as defined by ATP hydrolyzed per substrate transported, is more efficient for the disulfide linked variants relative to the wild-type *Na*Atm1. Since disulfide bond formation in these variants was not quantitative, we cannot eliminate some contribution of uncrosslinked material to the observed ATPase and transport activities, but the improved coupling efficiency relative to wildtype *Na*Atm1 revealed that the crosslinked protein is indeed functional with distinct properties.

**Table 2.** ATPase and transport activities for wildtype *Na* Atm1 and variants. (**a**) ATPase and transport activities with calculated coupling efficiencies of wildtype *Na*Atm1 and variants in units of nmol/min/mg. 10 mM MgATP and 2.5 mM GSSG were used in for these measurements. (**b**) The same ATPase and transport activities as tabulated in (**a**) in units of min^−1^. All activities were measured three times except for wildtype ATPPase activity, which was measured six times.

|  | *NaAtm1* | *NaA527C* | *NaS526C* | *NaT525C* |
| --- | --- | --- | --- | --- |
|  | ATPase activities (nmol/min/mg) |  |  |  |
| - GSSG (basal) | 64 ± 4 | 2.4 ± 0.6 | 54.0 ± 0.8 | 14.0 ± 0.7 |
| + GSSG (total) | 180 ± 15 | 6.5 ± 0.6 | 69.3 ± 0.7 | 20.8 ± 0.1 |
| Stimulated | 117 ± 16 | 4.1 ± 0.8 | 15 ± 1 | 6.8 ± 0.7 |
|  | Transport activities (nmol/min/mg) |  |  |  |
| Transport rate | 1.52 ± 0.02 | 0.16 ± 0.01 | 0.91 ± 0.08 | 0.50 ± 0.09 |
|  | Coupling efficiencies |  |  |  |
| Total/transport | 118 ± 10 | 41 ± 4 | 76 ± 6 | 42 ± 7 |
| Stimulated/transport | 77 ± 7 | 26 ± 6 | 17 ± 2 | 14 ± 3 |

**Table 2. ATPase and transport activities for wildtype *NaAtm1* and variants.**
|  | *NaAtm1* | *NaA527C* | *NaS526C* | *NaT525C* |
| --- | --- | --- | --- | --- |
|  | ATPase activities (min<sup>-1</sup>) |  |  |  |
| - GSSG (basal) | 8.4 ± 0.5 | 0.32 ± 0.07 | 7.2 ± 0.1 | 1.87 ± 0.09 |
| + GSSG (total) | 24 ± 2 | 0.86 ± 0.07 | 9.21 ± 0.09 | 2.77 ± 0.02 |
| Stimulated | 15.5 ± 0.3 | 0.5 ± 0.1 | 2.1 ± 0.1 | 0.90 ± 0.09 |
|  | Transport activities (min<sup>-1</sup>) |  |  |  |
| Transport rate | 0.203 ± 0.003 | 0.021 ± 0.001 | 0.12 ± 0.01 | 0.07 ± 0.01 |

An unanticipated finding of this work is that introduction of a disulfide bond between NBDs can improve the coupling efficiency. The disulfide crosslink between NBDs might be expected to inhibit transport by restricting relevant conformational changes and preventing full opening of the NBDs, as seen in the structural comparisons of the crosslinked chimeric ABCB1 in an occluded conformation (Alam et al. 2018) to the uncrosslinked inward-facing conformation of ABCB1 (Aller et al. 2009). The observation that covalently linked NBDs can transport substrate more efficiently, albeit at a somewhat reduced rate, suggests that the poor coupling typically observed for ABC transporters may reflect that the reconstituted systems are too dynamic or “floppy”. By restricting the conformational space through the introduction of disulfide crosslinks, the exploration of non-transport-relevant conformational states is apparently reduced, thereby improving the coupling efficiency.

The functional relevance of the uncoupled ATPase activity can be viewed from two perspectives that are not necessarily mutually exclusive. On the one hand, the uncoupled ATPase activity could reflect some aspect of the proteoliposome reconstitution system used to study ABC transporter function that does not faithfully mimic the native membrane, perhaps involving lipid composition. It is notable that significant variations in the coupling efficiency can be reported for a given transporter; for example, the coupling efficiency for the maltose transporter MalFGK in different studies ranges from 1.4 to over 3000 ATP/transported substrate (Table 1), which must arise from differences in the experimental protocols for reconstitution and transport characterization. On the other hand, the uncoupled activity could reflect the operation of two parallel, functionally relevant pathways for ATP hydrolysis that differ by the presence or absence of the transported substrate as proposed for certain ABC importers (Lewinson and Livnat-Levanon 2017) and ABC exporters (Hofmann et al. 2019). This behavior is in stark contrast to P-type transporters, a distinct class of ATP-dependent transporters where ATP hydrolysis is typically tightly coupled to transport (Palmgren and Nissen 2011). In this context, it may be relevant that P-type ATPases can exhibit turnover rates approaching 10^4^ min^−1^ (Skou 1998) which is considerably faster than the transport rates reported for ABC transporters (ranging between 1 to 1000 nmol/min/mg (Table 1), equivalent to 0.1 to 100 min^−1^ for a 100 kDa transporter). The lower turnover rates for ABC transporters would reduce the metabolic impact of uncoupled ATPase activity relative to P-type ATPases, which for the Na^+^, K^+^ ATPase represents a significant contribution to overall cellular energy consumption. The inefficient coupling of ABC transporters is also reminiscent of the properties of binding protein independent variants of the maltose transporter that exhibit constitutive ATPase activity (Covitz et al. 1994); this suggests the uncoupled ATPase activity may be associated with the ability to transport more weakly bound substrates. Understanding the origins of this uncoupling, and reducing the impact through disulfide crosslinking or other protocols, will be critical for the detailed dissection of the transport mechanism to assure that the ATP dependent steps are indeed relevant to substrate translocation.

## Author contributions

C.F. and D.C.R designed the research; C.F. performed the research; C.F., J.T.K and D.C.R. analyzed the data; and C.F and D.C.R prepared the manuscript.

## Data availability

Atomic coordinates were deposited in the Protein Data Bank with accession codes 6PAM (*Na*A527C-MgADP), 6PAN (*Na*S526C-ATP), 6PAO (*Na*T525C-ATP), 6PAQ (*Na*E523Q-ATP) and 6PAR (*Na*Atm1-MgAMPPNP). The raw data for ATPase and transport assays that support the findings in Figure 2 and Table 2 are included in Table S6, and all the other data are available from the corresponding author upon reasonable request.

## Acknowledgments

We thank the beamline staffs of the Stanford Synchrotron Radiation Lightsource beamline 12-2 and of the Advanced Photon Source GM/CA beamline for support during data collection. Discussions with Paul Adams, Gabriele Meloni, William Clemons, the organizers and speakers at the Cold Spring Harbor X-ray Method in Structural Biology Course (2018), the CCP4/APS School for Macromolecular Crystallography (2017) and the SBGrid/NE-CAT Phenix Workshop (2016) are gratefully acknowledged. We thank the Gordon and Betty Moore Foundation and the Beckman Institute for their generous support of the Molecular Observatory at Caltech. Use of the Stanford Synchrotron Radiation Lightsource, SLAC National Accelerator Laboratory, is supported by the U.S. Department of Energy, Office of Science, Office of Basic Energy Sciences under Contract No. DE-AC02-76SF00515. The SSRL Structural Molecular Biology Program is supported by the DOE Office of Biological and Environmental Research, and by the National Institutes of Health, National Institute of General Medical Sciences (including P41GM103393). GM/CA@APS has been funded in whole or in part with Federal funds from the National Cancer Institute (ACB-12002) and the National Institute of General Medical Sciences (AGM-12006). This research used resources of the Advanced Photon Source, a U.S. Department of Energy (DOE) Office of Science User Facility operated for the DOE Office of Science by Argonne National Laboratory under Contract No. DE-AC02-06CH11357. The Eiger 16M detector was funded by an NIH–Office of Research Infrastructure Programs, High-End Instrumentation Grant (1S10OD012289-01A1).

## Materials and Methods

### Mutagenesis and protein expression

The gene encoding *Na*Atm1 (with GenBank accession code ABD27067) was previously cloned into a pJL-H6 ligation independent vector with 6-His tag on the carboxy-terminus (Lee et al. 2014, Lee and Kim 2009), and deposited in Addgene (catalog #78308). All mutants were generated using Q5 Site-Directed Mutagenesis Kit (New England Biolabs). All constructs were overexpressed in *Escherichia coli* BL21-gold (DE3) cells (Agilent Technologies) using ZYM-5052 autoinduction media (Studier 2005). Selenomethionine substituted proteins were overexpressed in *Escherichia coli* B834 (DE3) cells (Novagen) using PASM-5052 autoinduction media (Studier 2005). All cells were collected by centrifugation and stored at −80 °C until use.

### Purification and crosslinking

Frozen cell pellets of cysteine mutants were resuspended in lysis buffer containing 100 mM NaCl, 20 mM Tris, pH 7.5, 40 mM imidazole, pH 7.5, 5 mM β-mercaptoethanol (BME), 10 mM MgCl_2_, 0.5% (w/v) n-dodecyl-β-D-maltopyranoside (DDM) (Anatrace), 0.5% (w/v) octaethylene glycol monododecyl ether (C12E8) (Anatrace), lysozyme, DNase, and protease inhibitor tablet. The resuspended cells were lysed either by solubilizing by stirring for 3 hours at 4 °C, or by using a M-110L pneumatic microfluidizer (Microfluidics). Unlysed cells and cell debris were removed by ultracentrifugation at 38,000 RPM for 45 minutes at 4 °C. The supernatant was collected and loaded onto a prewashed NiNTA column with NiNTA buffer A at 4 °C. NiNTA buffer A contains 100 mM NaCl, 20 mM Tris, pH 7.5, 50 mM imidazole, pH 7.5, 5 mM BME, 0.05% DDM and 0.05% C12E8. Elution was achieved using the same buffer with 350 mM imidazole. The eluted sample was then buffer exchanged to 100 mM NaCl, 20 mM Tris, pH 7.5, 0.05% DDM and 0.05% C12E8 (size exclusion chromatography (SEC) buffer). Oxidation of the introduced cysteines to form disulfide bonds was achieved by incubating buffer exchanged protein with 1 mM Cu (II)-(1,10-phenanthroline)_3_ for 1 hour at 4 °C. Crosslinked sample was then buffer exchanged into SEC buffer to remove the oxidant, and further purified by SEC on a HiLoad 16/60 Superdex 200 column (GE Healthcare). Fractions were pooled and concentrated using Amicon Ultra 15 concentrator (Millipore) with a molecular weight cutoff of 100 kDa to 20-35 mg/mL.

For cysteine-less constructs, the purified protein was prepared the same way as the cysteine constructs, but without BME. The eluted sample from NiNTA column was directly subjected to SEC without crosslinking or buffer exchange. Wildtype *Na*Atm1 was solubilized in lysis buffer containing 1% DDM and purified in NiNTA and SEC buffers containing 0.1% DDM for crystallization in the occluded conformation with MgAMPPNP.

### Proteoliposome preparation and transport assay

Proteoliposomes (PLS) were prepared by following published protocols for ABC transporters (Geertsma et al. 2008), with an additional step of Biobeads addition to ensure detergent removal. The transport assay was conducted with *Na*Atm1 reconstituted in PLS in a 1 mL format at 37 °C. The reaction mixture contained PLS at 5 mg/mL, 10 mM MgATP, pH 7.5, 2.5 mM GSSG, pH 7.5, and transport buffer at 85 mM NaCl, and 17 mM Tris, pH 7.5. The different controls were also prepared in similar fashion. 150 μL aliquots of the reaction mixture were taken every 15 minutes, added to 1 ml cold transport buffer, and then ultracentrifuged at 70,000 RPM in a TLA 100.3 rotor in a Beckman Ultima benchtop ultracentrifuge for 10 minutes at 4 °C. The pellets were washed 10 times with cold transport buffer, and then resuspended to 100 μl with solubilization buffer (85 mM NaCl, 17 mM Tris, pH 7.5 and 2% sodium dodecanoyl sarcosine (Anatrace)). The samples were solubilized for 2 hours until the solution clarified before spinning down in the TLA 100 rotor to remove bubbles. 10 μL samples were taken for GSSG quantification using the Glutathione Quantification Assay (Sigma-Aldrich). The rates were not corrected for orientation of *Na*Atm1 in PLS.

### ATPase assay

The ATPase activity was determined by the molybdate based phosphate quantification method (Chifflet et al. 1988). Briefly, all reactions were performed in a 250 μL scale with a final protein concentration of 0.05 mg/ml for both PLS and detergent at 37 °C. 50 μL of reactions were taken every 5 minutes for 4 times, mixed with 50 μL of 12% SDS in a 96-well plate at room temperature. 100 μL of ascorbic acid/molybdate mix was added, incubated for 5 minutes before the addition of 150 μL of citric acid/arsenite/acetic acid solution. The reaction was then incubated for 20 minutes at room temperature before reading at 850 nm with a Tecan plate reader. Reactions were done either in triplicates or sextuplicates, the absorbance measurements were plotted against time, and the final linear rates were fitted with nonlinear regression fit using Prism 8. The rates were not corrected for orientation of *Na*Atm1 in PLS.

### Crystallizations and structural determinations

*Na*A527C was crystallized in MemGold (Molecular Dimensions) condition #68. Upon optimization of the crystallization conditions, including additive screens (Hampton Research), the best crystals were grown from 100 mM NaCl, 100 mM Tris, pH 8.3, 25 mM MgCl_2_, and 28% polyethylene glycol 2,000 monomethyl ether (PEG 2000 MME) with 20 mM ATP at 20 °C. The *Na*A527C crystallization sample was prepared at 20 mg/mL with 1 mM ATP, 5 mM EDTA, with or without 5 mM GSSG. Crystals appeared in about 2 weeks and lasted for about 2 months. Crystals were harvested in cryoprotectant solutions containing 100 mM NaCl, 100 mM Tris, pH 8.3, 25 mM MgCl_2_, 28% PEG 2000 MME with PEG 400 at 10%, 15%, and 20% before flash-freezing in liquid nitrogen.

*Na*S526C, *Na*T525C and *Na*E523Q crystals were crystallized in the same condition as *Na*A527C except the removal of MgCl_2_ in the crystallization well solution. All crystallization samples were prepared with 1 mM ATP and 5 mM EDTA. The crystallization condition of *Na*T525C was further optimized with 200 mM of NDSB (non-detergents sulfobetaines)-221 using the Additive Screen (Hampton Research). The crystallization condition of *Na*E523Q was further optimized with 10 mM dithiothreitol but without 20 mM ATP in the crystallization conditions. Crystals of all three constructs were harvested in cryoprotectant solutions containing 100 mM NaCl, 100 mM Tris, pH 8.3, 28% PEG 2000 MME with PEG 400 at 10%, 15%, and 20% before flash-freezing in liquid nitrogen.

*Na*Atm1 purified in DDM was crystallized in MemChannel (Molecular Dimensions) condition #29. The crystallization sample was prepared in the presence of 1 mM AMPPNP, 2 mM MgCl_2_, and presence and absence of 5 mM GSSG. The condition was further optimized to 50 mM ADA, pH 7.1, 8-10% PEG 1000 and 8-10% PEG 1500 at 20°C with protein at 8 mg/ml. Crystals appears within a week. Crystals were harvested in cryoprotectant solutions containing 50 mM ADA, pH 7.1, 10% PEG 1000 and 10% PEG 1500 with PEG400 at 10%, 15%, and 20% before flash-freezing in liquid nitrogen.

### Data collection and structure determination

X-ray datasets were collected at the Stanford Synchrotron Radiation Laboratory beamline 12-2 using a Pilatus 6M detector with Blu-Ice interface (McPhillips et al. 2002) and the Advanced Photon Source GM/CA beamline 23ID-B using an Eiger 16M detector with JBluIce-EPICS interface (Stepanov et al. 2011). All datasets were processed and integrated with XDS (Kabsch 2010) and scaled with Aimless (Winn et al. 2011).

For the *Na*A527C crystal structure, the first 3 transporters in the asymmetric unit were identified by searching for multiple copies of the TMDs and NBDs using the original inward-facing structure (PDB ID: 4MRN) with Phaser in Phenix (Adams et al. 2010). Due to the relatively poor electron density for the fourth transporter, the helices of the TMDs were first built using Find Helices and Strands in Phenix (Adams et al. 2010), then a full transporter from the previously identified partial model was superposed onto the built helices in Coot (Emsley et al. 2010), which resulted in the misplacement of one NBD. The misplaced NBD was removed from the model and it was then correctly placed using Molrep in CCP4 (Winn et al. 2011). For the *Na*S526C structure, molecular replacement was carried out using Sav1866 (PDB ID: 2HYD) with superposed *Na*Atm1 sequence as the input model for Phaser in Phenix (Adams et al. 2010). For the *Na*T525C, *Na*E523Q and *Na*Atm1 fully occluded structures, molecular replacement was carried out using the *Na*S526C structure as the input model for Phaser in Phenix (Adams et al. 2010). For all structures, experimental phase information from SeMet datasets were obtained using with MR-SAD using AutoSol in Phenix (Adams et al. 2010). Iterative refinement and model building cycles were carried out with phenix.refine in Phenix (Adams et al. 2010), refmac in CCP4 (Winn et al. 2011) and Coot (Emsley et al. 2010), and the final refinements were carried out with phenix.refine in Phenix (Adams et al. 2010).

