## Supplementary figures for "Crosslinking of nucleotide binding domains improves the coupling efficiency of an ABC transporter"

**a**

```
                                   ABC signature                          Walker-B
Sav1866     459 FIMNLPQGYDTEVGERGVKLSGGQKQRLSIARIFLNNPP-------ILILDEATSALDLE 511
BtuD        114 ------LALDDKLGRSTNQLSGGEWQRVRLAAVVLQITPQANPAGQLLLLDEPMNSLDVA 167
NaAtm1      479 FIARLPQGYDTEVGERGLKLSGGEKQRVAIARTLVKNPP-------ILLFDEATSALDTR 531
                                                                    Targets for mutation
```

**b**

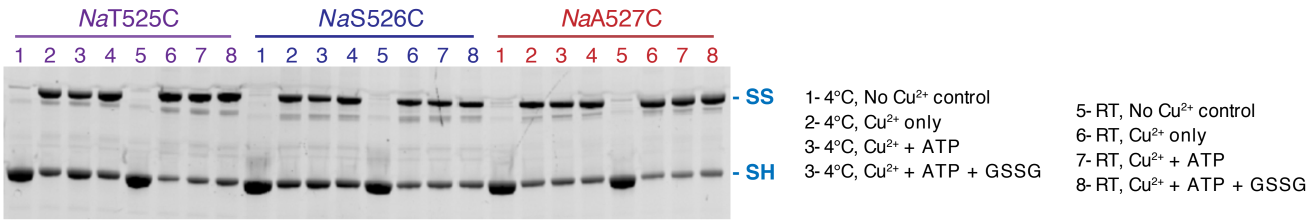


**Figure S1**. **Screening candidate residues in *Na*Atm1 for disulfide crosslinking**. (**a**) Partial sequence alignments of NBDs of various ABC transporters. (**b**)SDS-PAGE of the products of crosslinking with Cu (II)$_2$ (1,10-phenantroline)$_3$ under different conditions for the three cysteine variants in this report. RT= room temperature.

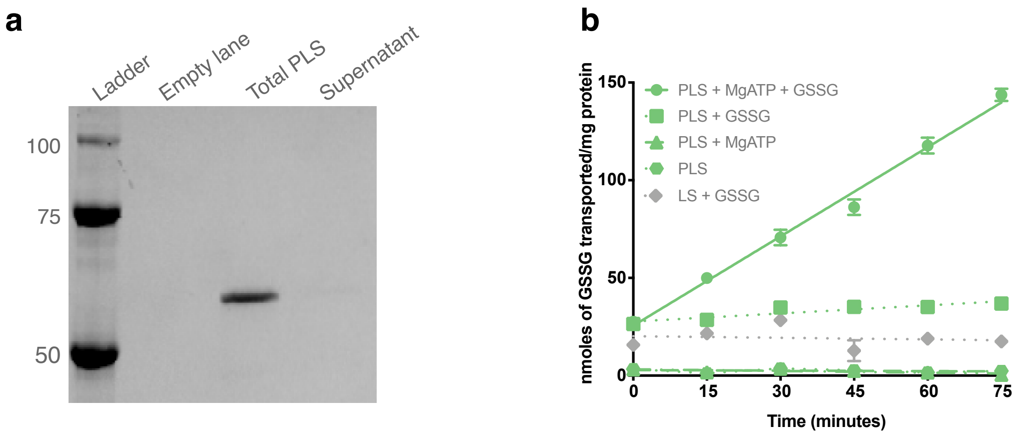


**Figure S2**. ***Na*Atm1 PLS reconstitution in PLS.** **a**) SDS-PAGE showing the reconstitution efficiency of *Na*Atm1 into PLS. **b**) Transport of GSSG by reconstituted *Na*Atm1 with various controls at 10 mM MgATP and 2.5 mM GSSG.

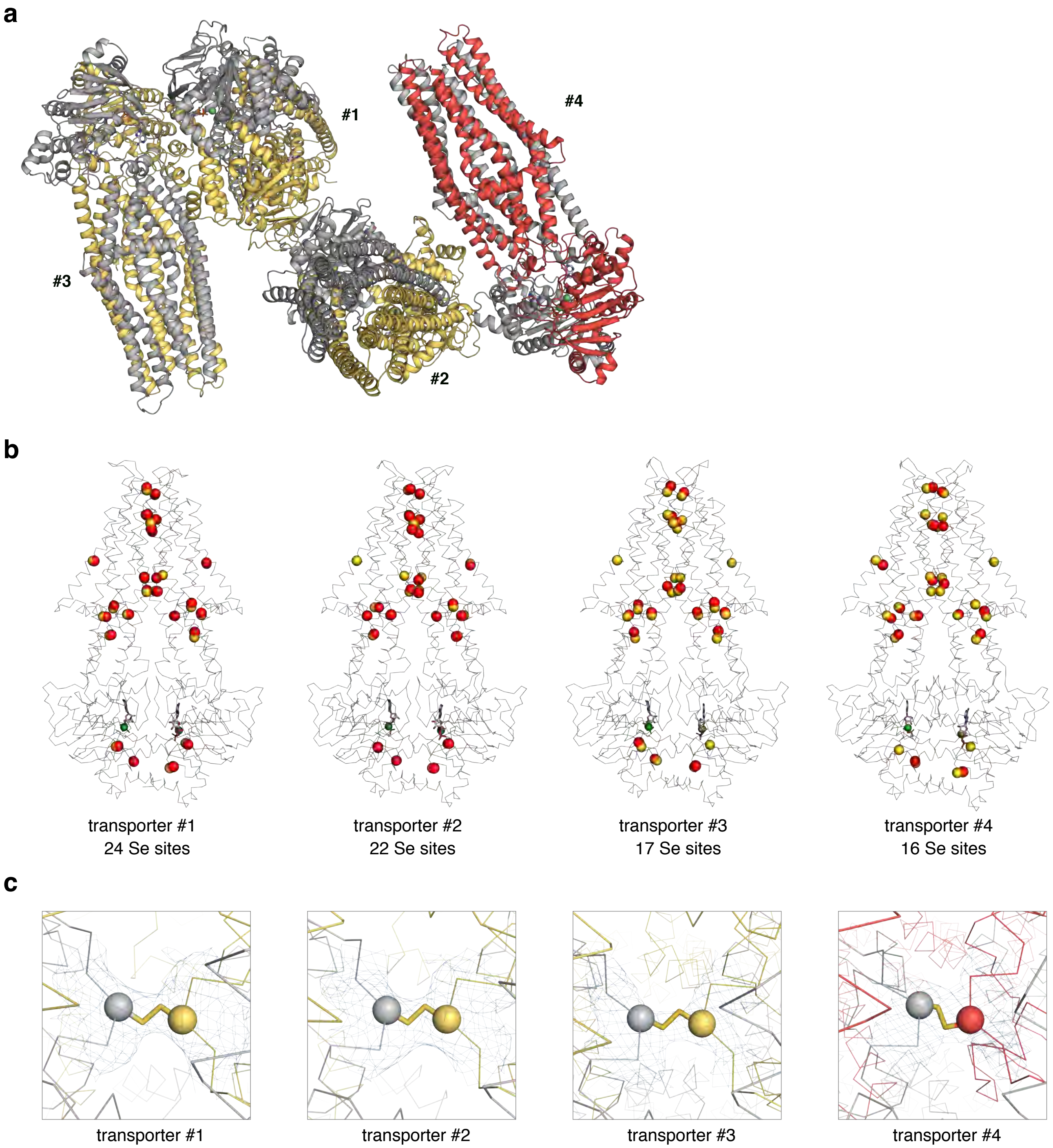


**Figure S3**. ***Na*A527C inward-facing occluded structures.** (**a**) The overall asymmetric unit of *Na*A527C in space group P1. (**b**) Location of selenium sites in the four transporters present in the selenomethionine-substituted *Na*A527C asymmetric unit. The selenium sites identified in Autosol in Phenix are shown in red spheres, the sulfur atoms of methionine residues from the refined model are shown in yellow spheres, ADP molecules are shown in sticks, and $Mg^{2+}$ ions are shown in green spheres. (**c**) Disulfide bridges in each of four transporters in the asymmetric unit. The C$\alpha$ carbons corresponding to C527 in the two chains are depicted as yellow and red spheres, respectively.

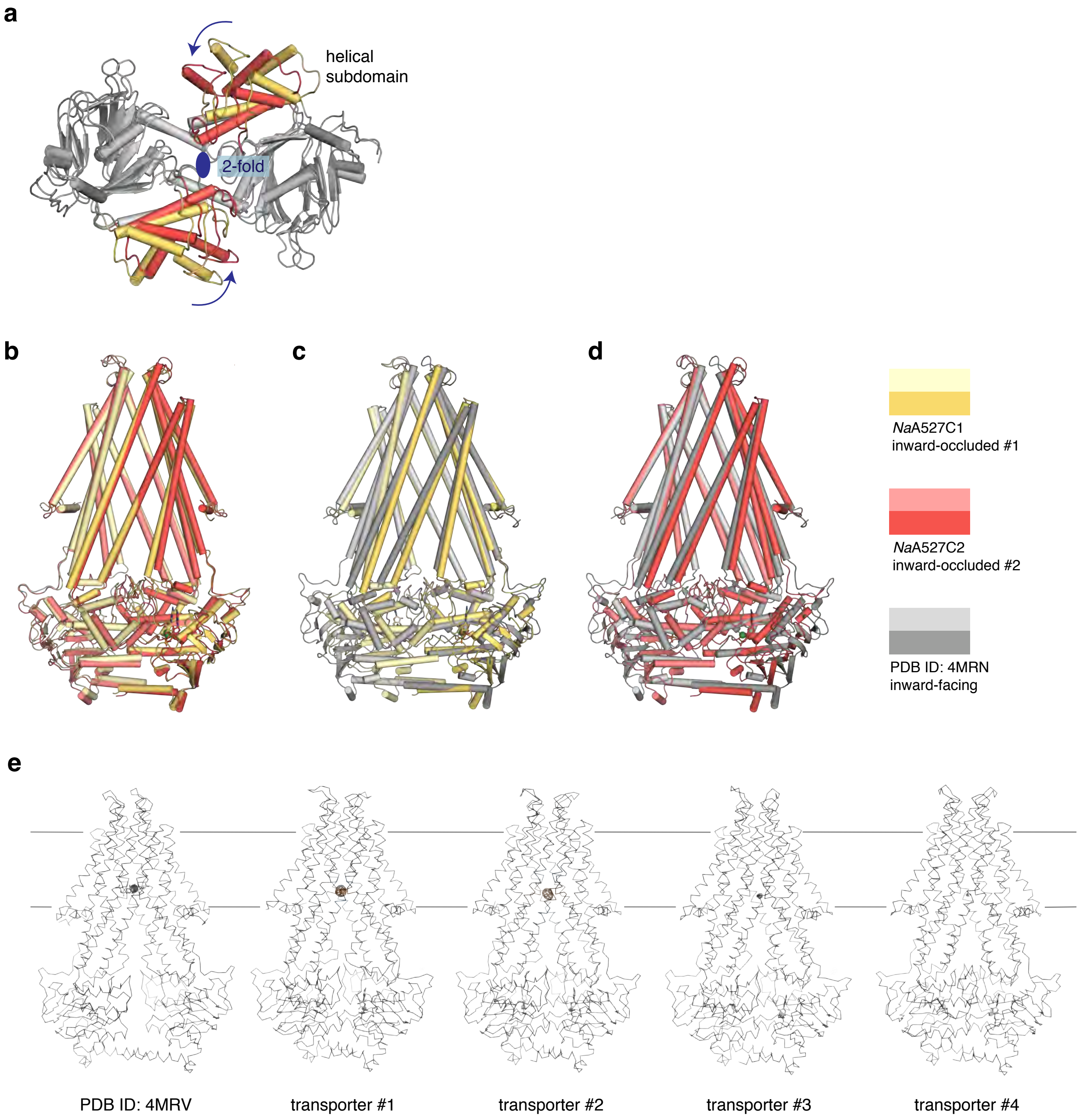


**Figure S4**. **Structural alignments of *Na*A527C in the inward-facing occluded conformations.** (**a**) Top-down view of the NBDs of the two *Na*A527C inward-facing occluded conformations after overall structural alignment. The molecular 2-fold axis is perpendicular to the screen. Arrows represent the relative rotations of the helical subdomains from inward-facing occluded #1 light grey to #2 in grey. The helical domains of inward-facing occluded #1 and #2 are colored in yellow and red, respectively. (**b**) Alignment of *Na*A527C inward-facing occluded #1 (paleyellow and yellow) to *Na*A527C inward-facing occluded #2 (light-red and red) with an overall RMSD of 1.7 Å. (**c**) Alignment of *Na*A527C inward-facing occluded #1 to *Na*Atm1 inward-facing conformation (lightgrey and grey) with an overall RMSD of 2.1 Å. (**d**) Alignment of *Na*A527C inward-facing occluded #2 to *Na*Atm1 inward-facing conformation with an overall RMSD of 4.4 Å. (**d**) Anomalous electron density map contouted at the 5 $\sigma$ levels (orange) for *Na*A527C crystallized in the presence of GS-Hg with comparison to the previous structure of *Na*Atm1 with GS-Hg bound (Hg shown in red sphere) (PDB ID: 4MRV).

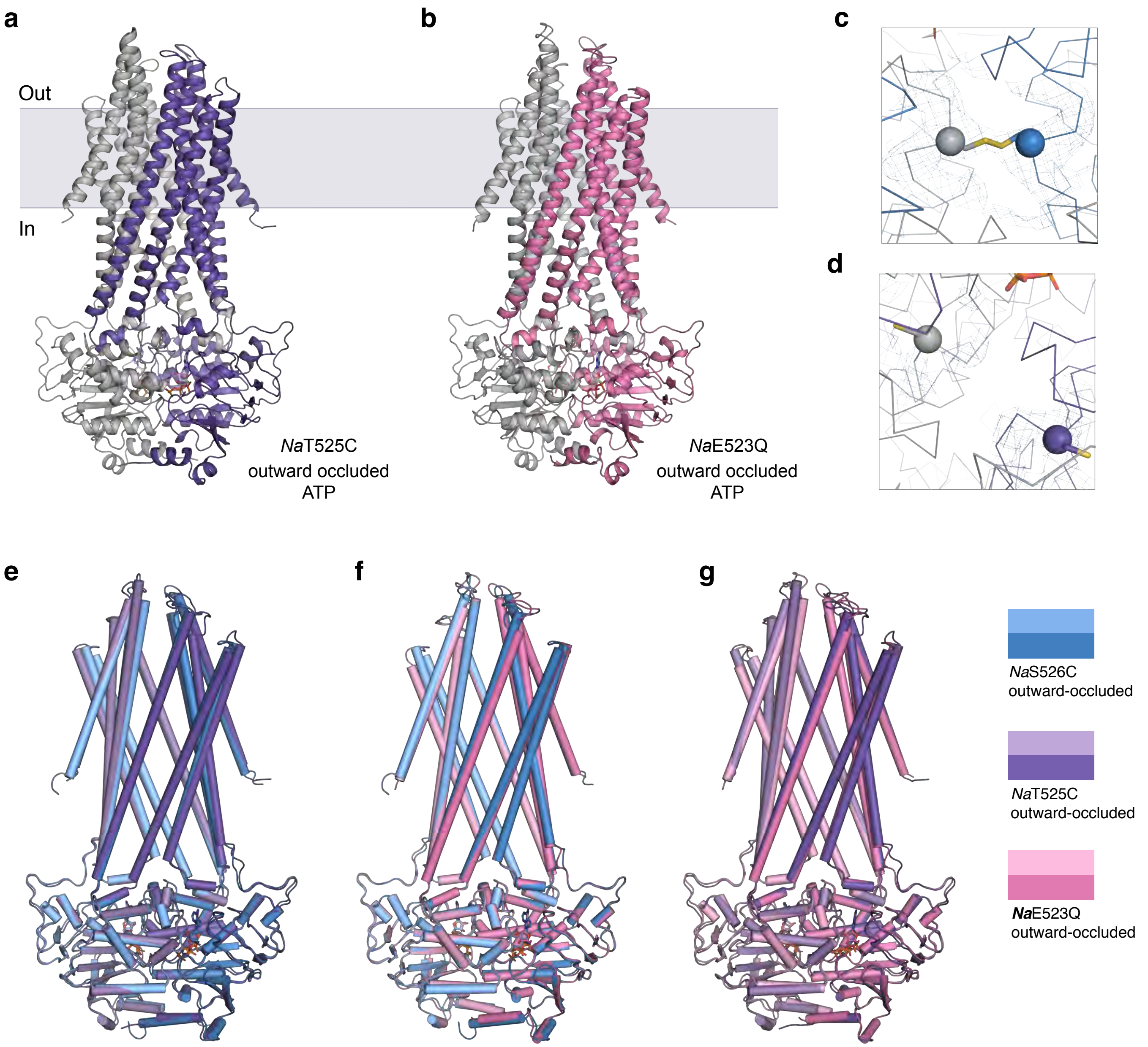


**Figure S5**. **Structural alignments of *Na*Atm1 in the outward-occluded conformations.** Crystal structures of *Na*T525C (**a**) and *Na*E523Q (**b**) in the outward-facing occluded conformations, both with ATP bound. (**c**) Disulfide bridge formed by S526C in the *Na*S526C structure with the C$\alpha$ in shown as grey and blue spheres. (**d**) T525C residues in the *Na*T525C outward-facing occluded structure with the C$\alpha$ positions shown as grey and purple spheres. (**e**) *Na*T525C overall structural alignment to *Na*S526C with an RMSD of 0.5 Å. (**f**) *Na*E523Q overall structural alignment to *Na*S526C with an RMSD of 0.5 Å. (**g**) *Na*E523Q overall structural alignment to *Na*T525C with an RMSD of 0.7 Å. *Na*S526C is colored in light blue and blue, *Na*T525C in light purple and purple, and *Na*E523Q in lightpink and pink.

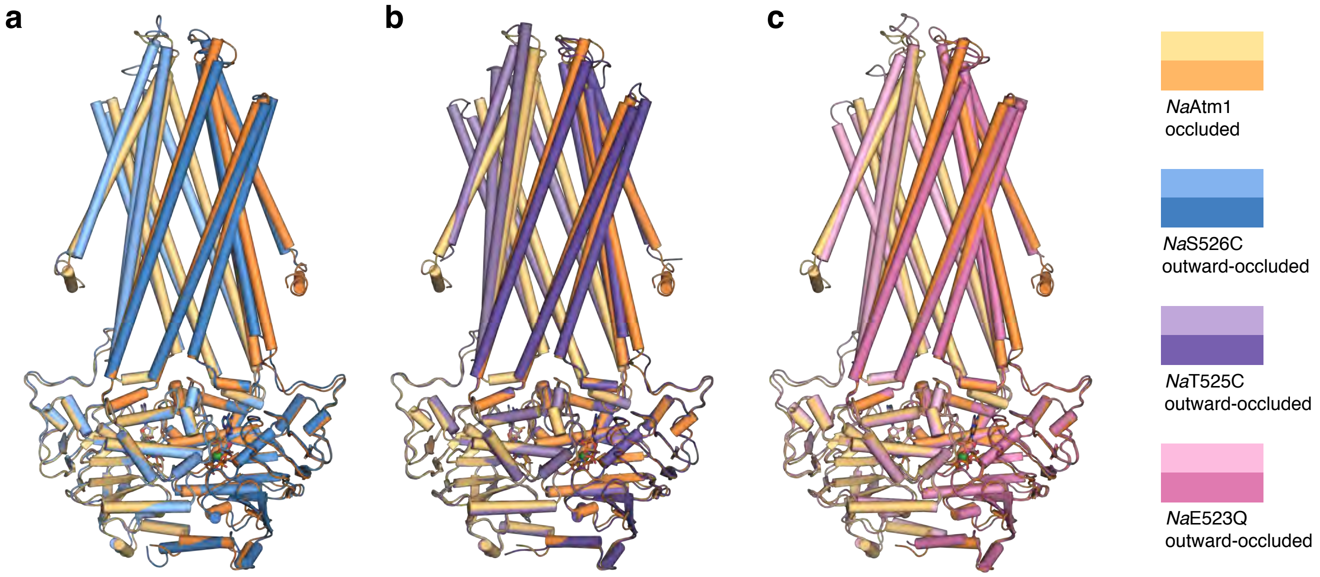


**Figure S6**. **Structural alignments of *Na*Atm1 in the fully occluded conformation.** *Na*Atm1 occluded structure alignments to (**a**) *Na*S526C with an overall RMSD of 1.1 Å, (**b**) *Na*T525C with an overall RMSD of 0.9 Å and (**c**) *Na*E523Q with an overall RMSD of 1.1 Å. *Na*Atm1 occluded structure colored in lightorange and orange, *Na*S526C in light blue and blue, *Na*T525C in light purple and purple and *Na*E523Q in lightpink and pink. AMPPNP is shown in sticks and $Mg^{2+}$ is shown in green spheres.
