## supplementary tables for "Crosslinking of nucleotide binding domains improves the coupling efficiency of an ABC transporter"

**Table S1. Data collection and refinement statistics of *Na*A527C**

| | ***Na*A527C native** | ***Na*A527C SeMet** |
|---|---|---|
| **Beamline** | SSRL 12-2 | SSRL 12-2 |
| **Wavelength (Å)** | 0.97946 | 0.97949 |
| **Resolution range (Å)** | 39.37 - 3.70 (3.832 - 3.70) | 39.72 - 4.50 (4.66 - 4.50) |
| **Space group** | P 1 | P 1 |
| **Unit cell (Å, °)** | 129.18 133.61 134.26 110.619 98.282 101.2 | 129.19 133.14 134.27 109.61 98.51 101.67 |
| **Total reflections** | 603652 (61595) | 972530 (99987) |
| **Unique reflections** | 84628 (7646) | 47277 (4667) |
| **Multiplicity** | 7.1 (7.3) | 20.6 (21.4) |
| **Completeness (%)** | 97.55 (89.28) | 99.3 (99.3) |
| **Mean I/sigma(I)** | 7.31 (0.74) | 8.1 (1.4) |
| **Wilson B-factor** | 141.79 | 163.64 |
| **R-merge** | 0.167 (3.050) | 0.291 (3.241) |
| **R-meas** | 0.180 (3.283) | 0.298 (3.320) |
| **R-pim** | 0.067 (1.206) | 0.066 (0.712) |
| **CC1/2** | 0.999 (0.412) | 0.998 (0.761) |
| **CC*** | 1.000 (0.764) | |
| **Reflections used in refinement** | 83473 (7643) | |
| **Reflections used for R-free** | 4185 (346) | |
| **R-work** | 0.240 (0.356) | |
| **R-free** | 0.288 (0.400) | |
| **CC(work)** | 0.787 (0.709) | |
| **CC(free)** | 0.841 (0.703) | |
| **Number of non-hydrogen atoms** | 36839 | |
| **macromolecules** | 36615 | |
| **ligands** | 224 | |
| **Protein residues** | 4722 | |
| **RMS (bonds) (Å)** | 0.004 | |
| **RMS (angles) (°)** | 0.95 | |
| **Ramachandran favored (%)** | 97.95 | |
| **Ramachandran allowed (%)** | 1.96 | |
| **Ramachandran outliers (%)** | 0.09 | |
| **Rotamer outliers (%)** | 0.40 | |
| **Clashscore** | 6.13 | |
| **Average B-factor** | 178.14 | |
| **macromolecules** | 178.04 | |
| **ligands** | 194.08 | |

**Statistics for the highest-resolution shell are shown in parentheses.

**Table S2. Data collection and refinement statistics of *Na*S526C**

| | ***Na*S526C SeMet** |
|---|---|
| **Beamline** | SSRL 12-2 |
| **Wavelength (Å)** | 0.97938 |
| **Resolution range (Å)** | 39.71 - 3.40 (3.522 - 3.40) |
| **Space group** | P 21 21 21 |
| **Unit cell (Å, º)** | 95.5 134.58 190.12 90 90 90 |
| **Total reflections** | 916579 (84633) |
| **Unique reflections** | 34343 (3016) |
| **Multiplicity** | 26.7 (25.1) |
| **Completeness (%)** | 98.64 (89.27) |
| **Mean I/sigma(I)** | 17.26 (1.19) |
| **Wilson B-factor** | 138.15 |
| **R-merge** | 0.148 (3.919) |
| **R-meas** | 0.151 (4.000) |
| **R-pim** | 0.029 (0.792) |
| **CC1/2** | 1.000 (0.588) |
| **CC*** | 1.000 (0.861) |
| **Reflections used in refinement** | 33962 (3012) |
| **Reflections used for R-free** | 1696 (153) |
| **R-work** | 0.192 (0.295) |
| **R-free** | 0.234 (0.377) |
| **CC(work)** | 0.727 (0.820) |
| **CC(free)** | 0.865 (0.750) |
| **Number of non-hydrogen atoms** | 8921 |
| **macromolecules** | 8859 |
| **ligands** | 62 |
| **Protein residues** | 1147 |
| **RMS (bonds) (Å)** | 0.002 |
| **RMS (angles) (º)** | 0.57 |
| **Ramachandran favored (%)** | 98.77 |
| **Ramachandran allowed (%)** | 1.23 |
| **Ramachandran outliers (%)** | 0.00 |
| **Rotamer outliers (%)** | 0.88 |
| **Clashscore** | 7.07 |
| **Average B-factor** | 196.93 |
| **macromolecules** | 196.93 |
| **ligands** | 197.86 |

**Statistics for the highest-resolution shell are shown in parentheses.

**Table S3. Data collection and refinement statistics of *Na*T525C**

| | ***Na*T525C native** | ***Na*T525C SeMet** |
|---|---|---|
| **Beamline** | APS GM/CA 23-IDB | APS GM/CA 23-IDB |
| **Wavelength (Å)** | 1.033202 | 0.979338 |
| **Resolution range (Å)** | 39.1 - 3.65 (3.78 - 3.65) | 39.31 - 3.90 (4.21 - 3.90) |
| **Space group** | P 21 21 21 | P 21 21 21 |
| **Unit cell (Å, º)** | 94.164 135.415 191.592 90 90 90 | 93.78 136.37 192.48 90 90 90 |
| **Total reflections** | 246061 (22321) | 237126 (49769) |
| **Unique reflections** | 27912 (2712) | 23170 (4675) |
| **Multiplicity** | 8.8 (8.1) | 10.2 (10.6) |
| **Completeness (%)** | 97.77 (97.54) | 99.9 (99.9) |
| **Mean I/sigma(I)** | 14.33 (1.11) | 12.1 (1.6) |
| **Wilson B-factor** | 172.32 | 180.32 |
| **R-merge** | 0.076 (2.002) | 0.100 (1.852) |
| **R-meas** | 0.080 (2.146) | 0.106 (1.945) |
| **R-pim** | 0.027 (0.758) | 0.033 (0.590) |
| **CC1/2** | 0.999 (0.634) | 1.000 (0.586) |
| **CC*** | 1.000 (0.881) | |
| **Reflections used in refinement** | 27348 (2699) | |
| **Reflections used for R-free** | 1356 (126) | |
| **R-work** | 0.251 (0.430) | |
| **R-free** | 0.285 (0.475) | |
| **CC(work)** | 0.713 (0.364) | |
| **CC(free)** | 0.877 (0.304) | |
| **Number of non-hydrogen atoms** | 8835 | |
| **macromolecules** | 8773 | |
| **ligands** | 62 | |
| **Protein residues** | 1135 | |
| **RMS (bonds) (Å)** | 0.003 | |
| **RMS (angles) (º)** | 0.63 | |
| **Ramachandran favored (%)** | 97.70 | |
| **Ramachandran allowed (%)** | 2.30 | |
| **Ramachandran outliers (%)** | 0.00 | |
| **Rotamer outliers (%)** | 0.22 | |
| **Clashscore** | 7.07 | |
| **Average B-factor** | 214.38 | |
| **macromolecules** | 214.56 | |
| **ligands** | 189.06 | |

**Statistics for the highest-resolution shell are shown in parentheses.

**Table S4. Data collection and refinement statistics of *Na*E523Q**

| | *Na*E523Q SeMet |
|---|---|
| **Beamline** | SSRL 12-2 |
| **Wavelength (Å)** | 0.97946 |
| **Resolution range (Å)** | 38.63 - 3.30 (3.419 - 3.30) |
| **Space group** | P 21 21 21 |
| **Unit cell (Å, º)** | 89.346 115.354 184.536 90 90 90 |
| **Total reflections** | 387847 (39220) |
| **Unique reflections** | 29277 (2861) |
| **Multiplicity** | 13.2 (13.6) |
| **Completeness (%)** | 99.32 (98.72) |
| **Mean I/sigma(I)** | 11.11 (1.01) |
| **Wilson B-factor** | 103.58 |
| **R-merge** | 0.190 (2.680) |
| **R-meas** | 0.200 (2.784) |
| **R-pim** | 0.054 (0.748) |
| **CC1/2** | 1.000 (0.700) |
| **CC*** | 1.000 (0.908) |
| **Reflections used in refinement** | 29179 (2845) |
| **Reflections used for R-free** | 1434 (127) |
| **R-work** | 0.234 (0.330) |
| **R-free** | 0.300 (0.439) |
| **CC(work)** | 0.684 (0.848) |
| **CC(free)** | 0.827 (0.615) |
| **Number of non-hydrogen atoms** | 8842 |
| **macromolecules** | 8780 |
| **ligands** | 62 |
| **Protein residues** | 1136 |
| **RMS (bonds) (Å)** | 0.002 |
| **RMS (angles) (º)** | 0.63 |
| **Ramachandran favored (%)** | 97.52 |
| **Ramachandran allowed (%)** | 2.48 |
| **Ramachandran outliers (%)** | 0.00 |
| **Rotamer outliers (%)** | 1.77 |
| **Clashscore** | 8.24 |
| **Average B-factor** | 134.03 |
| **macromolecules** | 133.82 |
| **ligands** | 163.57 |

**Statistics for the highest-resolution shell are shown in parentheses.

**Table S5. Data collection and refinement statistics of *Na*Atm1**

| | ***Na*Atm1 native** | ***Na*Atm1 SeMet** |
|---|---|---|
| **Beamline** | SSRL 12-2 | SSRL 12-2 |
| **Wavelength (Å)** | 0.97946 | 0.9793 |
| **Resolution range (Å)** | 39.33 - 3.35 (3.47 - 3.35) | 39.85 - 3.60 (3.67 - 3.60) |
| **Space group** | P 21 | P 21 |
| **Unit cell (Å, º)** | 169.648 92.498 237.691 90 110.34 90 | 170.10 92.21 237.47 90 110.58 90 |
| **Total reflections** | 686114 (60885) | 559063 (30390) |
| **Unique reflections** | 98507 (9041) | 79094 (4376) |
| **Multiplicity** | 7.0 (6.4) | 7.1 (6.9) |
| **Completeness (%)** | 97.85 (91.11) | 98.1 (95.8) |
| **Mean I/sigma(I)** | 9.43 (0.97) | 7.5 (1.1) |
| **Wilson B-factor** | 91.20 | 94.99 |
| **R-merge** | 0.174 (1.768) | 0.211 (1.942) |
| **R-meas** | 0.189 (1.927) | 0.228 (2.100) |
| **R-pim** | 0.071 (0.754) | 0.086 (0.787) |
| **CC1/2** | 0.999 (0.530) | 0.994 (0.466) |
| **CC*** | 1.000 (0.832) | |
| **Reflections used in refinement** | 97914 (9035) | |
| **Reflections used for R-free** | 4897 (425) | |
| **R-work** | 0.254 (0.353) | |
| **R-free** | 0.282 (0.365) | |
| **CC(work)** | 0.740 (0.685) | |
| **CC(free)** | 0.529 (0.606) | |
| **Number of non-hydrogen atoms** | 26975 | |
| **macromolecules** | 26783 | |
| **ligands** | 192 | |
| **Protein residues** | 3464 | |
| **RMS (bonds) (Å)** | 0.003 | |
| **RMS (angles) (º)** | 0.60 | |
| **Ramachandran favored (%)** | 97.35 | |
| **Ramachandran allowed (%)** | 2.59 | |
| **Ramachandran outliers (%)** | 0.06 | |
| **Rotamer outliers (%)** | 0.58 | |
| **Clashscore** | 8.33 | |
| **Average B-factor** | 120.02 | |
| **macromolecules** | 120.15 | |
| **ligands** | 101.67 | |

**Statistics for the highest-resolution shell are shown in parentheses.

### Table S6a. Raw transport activities for all variants

| Samples | Time (min) | | | | | |
|---|---|---|---|---|---|---|
| | 0 | 15 | 30 | 45 | 60 | 75 |
| ***Na*****Atm1 PLS +MgATP +GSSG** | 28.17 | 45.49 | 69.95 | 81.08 | 113.06 | 141.67 |
| | 28.05 | 49.95 | 64.09 | 94.01 | 125.89 | 139.62 |
| | 26.57 | 54.53 | 77.96 | 83.45 | 114.33 | 149.99 |
| ***Na*****Atm1 PLS +GSSG** | 26.69 | 26.22 | 30.82 | 34.07 | 34.73 | 39.73 |
| | 26.08 | 28.78 | 37.44 | 36.06 | 38.74 | 32.31 |
| | 26.59 | 30.69 | 36.27 | 35.48 | 31.99 | 38.69 |
| ***Na*****Atm1 PLS +MgATP** | 2.84 | 1.03 | 2.95 | 2.98 | 1.13 | 1.48 |
| | 3.39 | 1.59 | 3.58 | 2.29 | 2.17 | 1.83 |
| | 3.39 | 2.54 | 4.10 | 2.08 | 1.65 | -1.82 |
| ***Na*****Atm1 PLS** | 3.30 | 1.80 | 3.34 | 2.64 | 1.48 | 1.76 |
| | 3.39 | 1.76 | 3.75 | 2.18 | 1.31 | 1.59 |
| | 3.39 | 0.16 | 4.03 | 3.30 | 2.04 | 3.95 |
| **LS +GSSG** | 20.25 | 22.67 | 29.57 | 17.30 | 20.47 | 16.88 |
| | 14.00 | 21.70 | 25.88 | 18.91 | 16.76 | 16.41 |
| | 12.87 | 20.42 | 29.74 | 2.07 | 19.66 | 19.29 |
| ***Na*****A527C PLS +MgATP +GSSG** | 23.88 | 19.64 | 31.19 | 30.18 | 26.25 | 37.11 |
| | 24.36 | 26.97 | 32.05 | 31.50 | 33.88 | 35.79 |
| | 19.28 | 26.69 | 33.64 | 31.78 | 29.64 | 35.41 |
| ***Na*****S526C PLS +MgATP +GSSG** | 22.86 | 42.06 | 53.21 | 66.26 | 77.55 | 79.59 |
| | 21.26 | 41.68 | 59.75 | 61.69 | 84.84 | 92.92 |
| | 22.58 | 37.91 | 61.09 | 64.64 | 82.63 | 103.12 |
| ***Na*****T525C PLS +MgATP +GSSG** | 23.32 | 25.75 | 15.61 | 49.92 | 58.39 | 63.36 |
| | 20.60 | 26.69 | 40.53 | 51.72 | 52.88 | 37.39 |
| | 21.82 | 25.94 | 39.12 | 42.16 | 59.62 | 57.18 |
| ***Na*****E523Q PLS +MgATP +GSSG** | 21.82 | 26.31 | 30.16 | 31.69 | 32.19 | 32.12 |
| | 16.01 | 24.62 | 30.91 | 8.99 | 35.00 | 33.71 |
| | 17.88 | 21.80 | 23.30 | 32.07 | 35.00 | 34.47 |

**Table S6b. Raw ATPase activities for all variants**

| Conditions | Variants | | | | |
|---|---|---|---|---|---|
| | *Na*Atm1 | *Na*A527C | *Na*S526C | *Na*T525C | *Na*E523Q |
| Proteoliposomes + 10mM MgATP | 51.90 | 1.26 | 55.21 | 13.28 | 1.22 |
| | 77.01 | 2.97 | 52.58 | 15.39 | 1.86 |
| | 67.60 | 2.93 | 54.32 | 13.43 | 1.00 |
| | 56.09 | | | | |
| | 68.27 | | | | |
| | 60.02 | | | | |
| Proteoliposomes + 10mM MgATP + 2.5mM GSSG | 118.68 | 7.55 | 69.50 | 20.88 | -0.08 |
| | 179.94 | 5.66 | 70.29 | 21.00 | -0.46 |
| | 157.94 | 6.18 | 68.06 | 20.55 | 0.48 |
| | 216.80 | | | | |
| | 206.72 | | | | |
| | 201.87 | | | | |
| Detergent (DDM/C12E8) + 10mM MgATP | 122.53 | 11.50 | 108.90 | 57.47 | 5.20 |
| | 125.12 | 13.32 | 111.60 | 57.37 | 5.76 |
| | 97.76 | 12.30 | 112.40 | 56.94 | 3.30 |
| | 134.40 | | | | |
| | 127.08 | | | | |
| | 131.24 | | | | |
| Detergent (DDM/C12E8) + 10mM MgATP + 2.5mM GSSG | 215.77 | 14.16 | 104.30 | 42.67 | 3.25 |
| | 200.91 | 15.01 | 103.80 | 42.01 | 4.76 |
| | 202.41 | 15.10 | 103.20 | 42.88 | 4.26 |
| | 288.59 | | | | |
| | 290.64 | | | | |
| | 295.58 | | | | |
